# Regulating AMPKα and insulin level by Vinegar, Swimming and Refeeding on High-Fat Diet Rats to Rebuild Lipid Homeostasis

**DOI:** 10.1101/2020.01.08.899419

**Authors:** Yuan Yang, Feng Zhang, Xiao Xiao, Chunlian Ma, Hua Liu, Yi Yang

## Abstract

Our aims were to explore the effects of dietary and behavior interventions on lipometabolism caused by unhealthy high-fat diet and the best method to rebuild lipid homeostasis of this lifestyle. Apart from normal diet rats, 34 rats were fed with high-fat emulsion for 4 weeks before being divided into 4 groups and intervened for another 4 weeks. 8 of them were classified into high-fat control group and 9 were sorted into high-fat diet with rice vinegar group. Meanwhile, 10 were put into high-fat diet with swimming group and 7 were just for refeeding normal diet group. Then the data of body weight was recorded and analyzed. Serum, pancreas, liver, cardiac tissues and epididymis adipose were sampled as required. Indexes of serum were tested by kits. AMPKα, HNF1α, CTRP6 from tissues were detected by western blot. According to our experiments, Swimming and refeeding groups reflected a better regulation on lipid homeostasis mainly by up-regulating the expression of pancreas AMPKα. To be more specific, the refeeding rats showed lower T-CHO (*P*<0.001) and LDL-C (*P*<0.05), but higher weight gain (*P*<0.001),insulin level (*P*<0.01)and pancreas AMPKα (*P*<0.01)than high-fat control rats. Compared with rats experimented by swimming or rice vinegar, they showed higher weight gain (*P*<0.001),insulin level (*P*<0.01)and HNF1α, but lower of CTRP6. In summary, refeeding diet functioned better in regulating the lipometabolic level after high-fat diet. Whatever approach mentioned above we adopted to intervene, the best policy to keep the balance of lipid homeostasis is to maintain a healthy diet.

## 3. Introduction

High-fat diet is the main cause of abnormal lipometabolism, such as Hyperlipidemia, Obesity and Cardiovascular disease. It is likely to be followed by unstable insulin level, imbalanced energy metabolism and depraved dyslipidaemia (1). Although the content of plasma lipids counts small part of whole-body lipid, both exogenous and endogenous lipids need to be run among tissues through the blood. Therefore, changes in blood lipids can basically reflect the status of lipid metabolism in the body. That is, regulating the balance of lipid intake and exclusion is essential to maintain lipid metabolism. Excessive intake of lipids from diet affects flux of substrates through lipogenesis and lipid oxidation to intervene related metabolism and insulin biological function (2). Since abnormal lipometabolism tends to develop into chronic refractory diseases with multi-tissues and organs (3), superior methods should be remarked as long-term and progressive effects. At present, some studies have shed light on the potential lipid-lowering properties by some certain ways, e.g. swimming (4, 5), functional foods and dietary supplements(e.g. rice vinegar) (6, 7). They may restore the balance of lipid metabolism by enhancing lipid metabolism and increasing exclusion. However, these efficacious measures with quantitative standard may not be suitable for individuals to practice. As reducing lipid intake is the other way to regulate the imbalanced metabolism, low-fat diet and ketogenic diet are also heatedly discussed among the general diet modes (8, 9). Here, we hypothesize that refeeding diet, an acceptable diet of returning to normal diet from high-fat dietary condition, can also result in lowering lipid by certain ways (Fig.1).

**Fig1.**
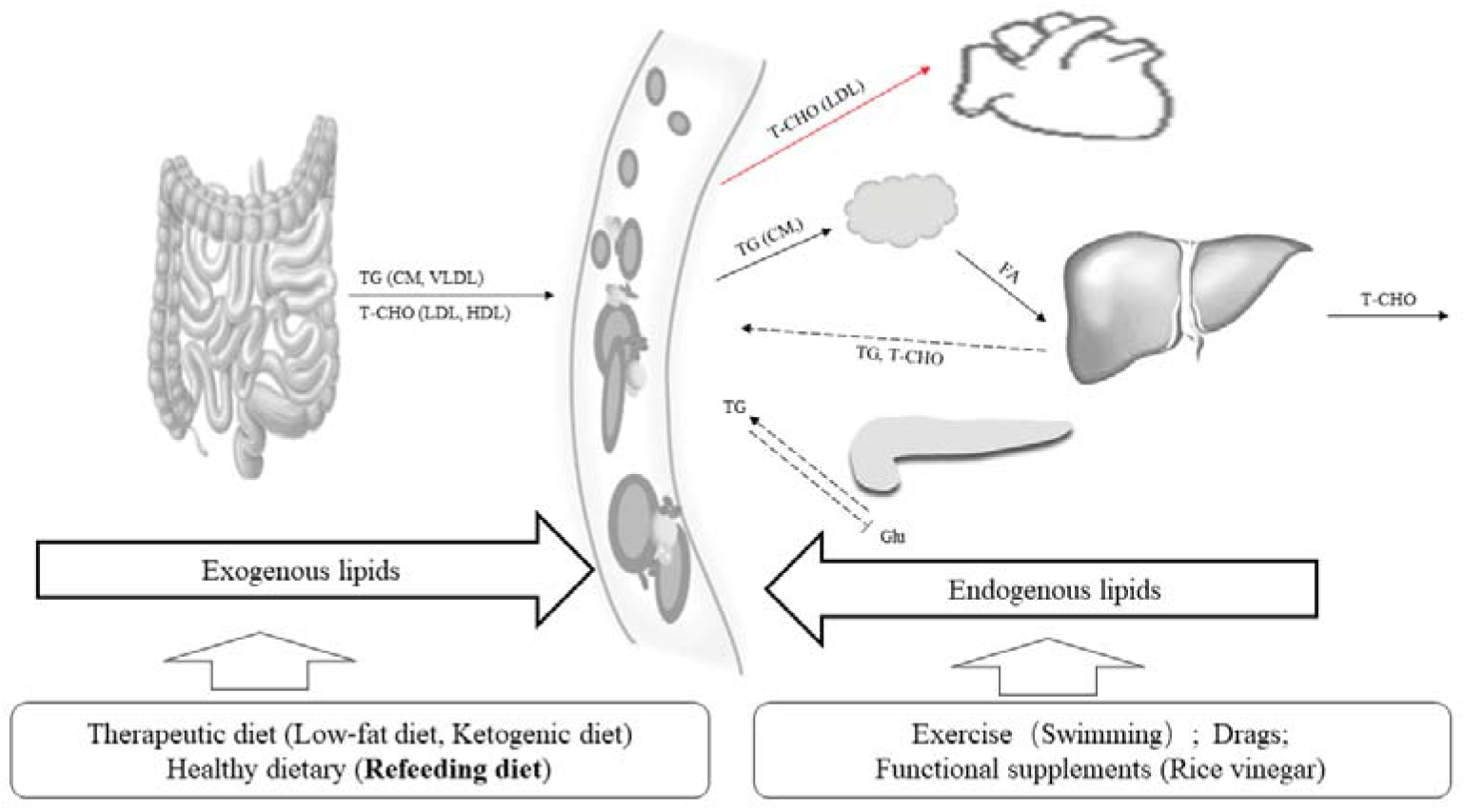
Balance of blood lipid metabolism and corresponding interventions. Exogenous and endogenous lipids maintain blood lipid levels. Excessive fat intake needs to accelerate the metabolism and limit the synthetic lipids in vivo to restore the balance of blood lipids. The process involves the coordination of multiple organs: ①Excessive fatty acids (FA) in adipose tissue need to be metabolized through the liver and bile; ②Reducing insulin-mediated glucide (Glu) converses to excessive TG; ③Reducing the synthesis of TG and T-CHO in liver. Excessive T-CHO will come to the heart from blood circulation and cause serious cardiovascular diseases. There are two ways to regulate the abnormal lipid metabolism. One is a therapeutic diet that blocking dyslipidemia from exogenous sources. The other way is to exert biological effects on the corresponding pathway and regulate lipid metabolism in vivo by exercise, drugs, functional supplements and so on.

As increasing evidence shows, AMP-activated protein kinase (AMPK) has emerged as an attractive target sensor to modulate lipid abnormalities and maintain energy homeostasis(10). The strategy related to AMPK pathway on the regulation of whole-body lipid metabolism has been of interest, especially on the higher effect of heart, liver, and gonadal adipose tissue as compared with skeletal muscle by AMPK (11). Moreover, Liver and peripheral fat are important insulin-targeted tissues which can reflect global lipometabolic level (21). Here we focused on the related AMPK substrates: C1q/TNF-associated protein (CTRP) and hepatocyte nuclear factor-1 α (HNF1α). CTRP6 is an essentially insulin-sensitive regulator to link adipose tissue and insulin resistance by modulating lipid development and metabolism in peripheral tissues (12). HNF1α, the typical transcription factor in liver, is involved in regulating hepatocyte lipometabolism and pancreas-induced insulin level (13). In order to reveal the best way to regulate lipometabolism by refeeding diet, rice vinegar or swimming, the lipometabolism related AMPKα in pancreas, liver and cardiac tissues of high-fat diet rats were observed. The regulation of AMPK/CTRP6 and AMPK/HNF1α on fatty acid metabolism specifically in peripheral and liver tissues were also analyzed. Our ultimate goal is to explore the best lipid-lowering way to cope with abnormal lipometabolism problem, and to provide a more scientific theoretical basis for a healthier dietary lifestyle finally.

## 4. Materials and Methods

### Materials

#### Ethics

All animal experimental protocols in the study were reviewed and approved by the Institutional Animal care and Use committee of Wuhan Sports University. They were carried out under the guidelines of the National institute of Health Guide for the Care and Use of Laboratory Animals.

#### Animals

39 SPF male Sprague-Dawley rats aged 2 months with the weight between 200±15 g were studied. These rats were purchased from the Research Center of Laboratory Animal in Hubei Province (No.42000600014140). They were housed individually in stainless steel cages under controlled conditions (22±3 °C, 40%~70% humidity and 12:12-h light-dark cycle) for 1 week‵s adjustment and then incorporated into our experiment. Apart from normal diet group (NC, n=5), other rats were fed with high-fat emulsion for 4 weeks and intervened by different methods for following 4 weeks including high-fat control group (HC, n=8), high-fat diet with rice vinegar group (HV, n=9), high-fat diet with swimming group (HS, n=10), and refeeding normal diet group (RH, n=7). All rats were given free diet and kept in the clean SPF animal lab to ensure the experimental research was not affect by various factors. In addition, we observed and recorded the rats’ spirit, diet, agility, defecation, hair and color daily.

#### Reagents& Equipment

Most reagents for SDS-PAGE and Oil Red O staining were purchased from Amresco or Biyuntian Bio of Shanghai. Others were listed as follows: HDL-C/LDL-C/TG/T-CHO test kit (Jiancheng Bio, Nanjing); Rat insulin ELISA kit (Huamei Bio, Wuhan); Rabbit anti CTRP6 (Boosen Bio, Beijing); Rabbit anti rat AMPKα (Emijie, Wuhan); Electrothermal thermostat incubator (Yuejin Medical, Shanghai); Vortex oscillator (Zhongxi Yuanda, Beijing); Multifunctional enzyme Standard instrument (Thermo Fisher, China); Centrifuge (Auchuangye, Beijing); Constant temperature culture oscillator (Zhicheng, Shanghai); Organization grinder (Jingxin, Shanghai); Ultrasonic cell crusher (Xinzhi Bio, Ningbo); Super-thermostat water bath pot/Magnetic agitator (Guohua Electric, Changzhou); Electrometer/Vertical electrophoresis bath (Liuyi, Beijing); Horizontal Shaker (Qilinbeier, Jiangsu); BX53 Biomicroscopy (Olympus, Beijing).

### Methodology

#### Diet protocol

After adaptive feeding, rats from HC, HS, HV and RH in the first 4 weeks were taken high-fat emulsion(20 g of lard, 2 g of Cholesterol, 2 g of Sodium Cholate, 20 ml of Tween 80, 20 ml of Propylene Glycol and 1 g of Propylthiouracil) (14–16). These high-fat diet rats were given intragastric administration of 5 ml/kg high-fat emulsion daily (17), and normal diet rats were given the same amount of distilled water every day. Rice vinegar of 9 degree was diluted into 3 degree and stored in the refrigerator. Each rat from HV was given 10 ml/kg of rice vinegar daily for 4 weeks.(18, 19)

#### Exercise protocol

Our exercise protocol was followed by the previous research of aerobic exercise for reducing animal stress and promoting health (20). Only one group member of our research was responsible for the animal management and intervention on the rats of HS group to avoid personality error. We adopted a moderate protocol by 60-minute swimming for 4 weeks (7 times/week) on average (21, 22). These rats underwent physical training within tepid water (30±2 °C) in experimental swimming pools (36 cm of depth).

BW (body weight, g) of each rat were measured and recorded weekly throughout the experiment for further analysis. After 4 weeks of intervention, the rats were anesthetized with weight-adjusted injection of 1% sodium pentobarbital (4 ml/kg) followed by 16 h of fasting and inactivity. Thoracic blood was collected and used to get the serum which was stored at −20 °C for the determination of metabolite concentrations. The adipose tissue samples (5 mm × 5 mm × 20 mm) in epididymis were quickly dissected in liquid nitrogen, then transferred to −20 °C refrigerator for 30min and reserved for frozen tissue slices. The liver, pancreas and epididymal fat tissues were quickly collected and snapped frozen in liquid nitrogen, and stored at −80 °C.

#### Histological analysis

Epididymis fat tissues were carefully dissected and placed on cork with oxytetracycline. These fragments were cut into 5~10 um thick and covered on sliders which were dried within 60 min at room temperature and then fixed in cold 10% formalin for 10 min. Then the slides were rinsed immediately with distilled water, soaked in 60% of isopropanol, and stained with Oil Red O solution for 10 min. After separated with 60% of isopropyl alcohol until the background was colorless, the slides were washed again and stained with Mayer hematoxylin, and then blued and distilled for several minutes. Finally, epididymis adipose slides were sealed and analyzed to get results of the degeneration and other changes of them.

#### Enzyme assay

In order to collect supernatant for serum tests, the blood stored at −20 °C was transferred to the freezer at 4°C for 30 min and centrifuged in centrifuge for 10 min the next day. Analysis of metabolite concentrations (total cholesterol (T-CHO), triacylglycerol (TG), cholesterol fractions (LDL-C/HDL-C) and insulin) were determined by enzymatic commercial kits followed by manufacturer’s instructions. The test methods of T-CHO and TG in rat serum were the same which directly determined by single reagent COD-PAP method according to the kit. Serum LDL-C and HDL-C were directly detected by the method according to the kit instructions. The content of insulin in serum was detected by ELISA method according to the manufacturer’s instructions. The main steps of these indexes above were to add buffer, sample and reaction solution with different concentration gradient in turn, to measure the OD value at different wavelengths, and to calculate the corresponding index concentration according to the formula for analyzing the content of serum samples. Samples from one experiment were running on the same plate in duplicate. And the average coefficient of variation was<10%.

#### Western blot

The expressions of AMPKα (in pancreas, cardiac and liver tissues), HNF1α (in liver) and CTRP6 (in epididymis adipose) were detected by protein immunoblotting. Samples were homogenized on ice with Organization grinder and Ultrasonic cell crusher (speed 10, 5×3s pulses) in RIPA buffer (ratio ~1:40), supplemented with protease and phosphatase inhibitors, and then centrifuged at 4°C (15 min with 12,000 rpm). The collected supernatant was used to determine protein content by a bicinchoninic acid assay. These protein samples, 10~20 ug of protein, were prepared in loading buffer (ratio 1:1) and separated by electrophoresis on 10% SDS-PAGE gels (45 V, 45 min; 85 V, 90 min). Proteins were wet transferred onto nitrocellulose membranes at 350 mA with 2 h. Membranes were shaken in blocking buffer prepared with 5% nonfat dry milk in Tris-buffered saline/0.1% Tween 20 (TBST) for 1 h. Then coming to the process of overnight incubation at 4 °C with the primary antibody (dissolved in blocking buffer with 1:1000). Following primary incubation, membranes were rinsed with TBST and incubated with horseradish peroxidase-conjugated secondary antibodies (dissolved with 1:10000) for 1 h at room temperature. Finally, signals were detected by enhanced chemiluminescence and subsequently quantified by densitometry and analyzed by Image J software. The proteins expressed by the ratio of gray value of protein band were normalized to β-actin.

#### Statistical analysis

The measured data were analyzed and processed by Spss19.0, GraphPad prism5, Image-pro plus 6.0 and CurveExpert1.3. Comparisons between different groups were made using unpaired, two-tailed t-tests. Body weight of different groups over time was determined using a one-way ANOVA followed by Tukey’s post hoc analysis. The data analyzed were expressed as Mean ± SE, *P*<0.05 indicated statistical difference, *P*<0.01 indicated that each data had significant difference, *P*<0.001 indicated extremely significant difference.

## 5. Results

### 1 Body Weight Changes by High-Fat Diet and Different Interventions

The analysis of body weight changes of two experimental periods (1~5 week and 5~9 week) were taken from a weekly-statistic protocol. In Fig.2(A), an upward trend of increase in the weight of rats was noticed in the first 4 weeks while statistic difference didn’t become obvious until the following four weeks. Combined with Fig.2(B), except for the consistently increasing trend in normal diet group and the refeeding group, the body weight of other three groups were decreasing distinctly. Specifically, the body weight gain of HC, HV and HS rats during 5~9 weeks was significantly lower than that of other two groups (NC and RH) (*P*<0.001) in Tab.1. However, there was no difference among HC, HV and HS groups, nor significantly difference on the body weight gain of RH and NC during 5~9 weeks. In brief, the weight of rats from refeeding diet kept increasing, while the continuous high-fat diet with or without swimming and vinegar experienced weight loss.

**Fig2.**
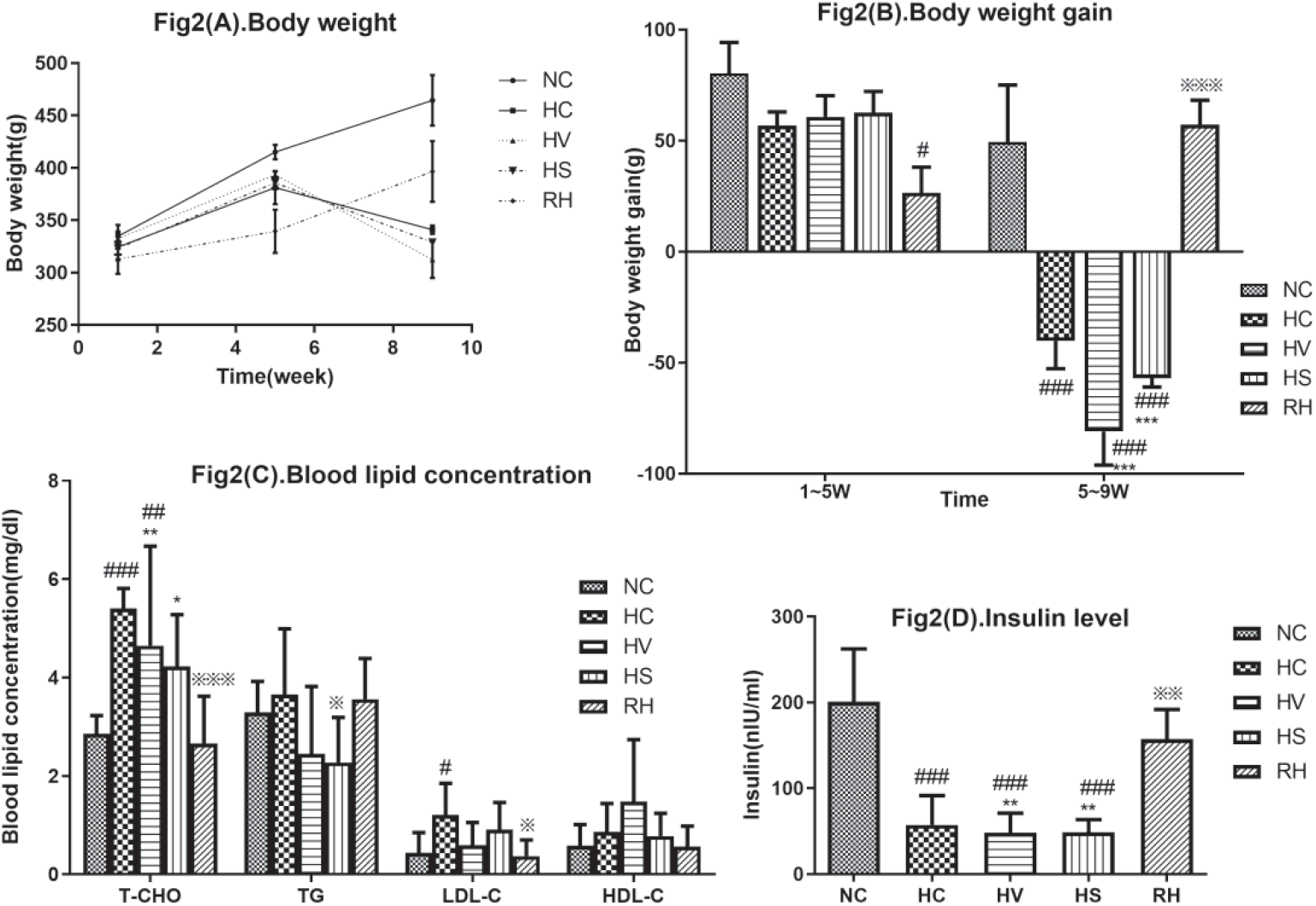
Weight and Blood analysis. Values are means ± SE; Compared with NC, #*P*<0.05, ^###^ *P*<0.001; Compared with HC, ^※^*P*<0.05, ^※※※^*P*<0.001; Compared with RH*, \**P*<0.05, \*\*\**P*<0.001; (A) and (B): note that, 0~1w indicated the weight of rats fed adaptively for 1 week with no significant body weight changing among different groups, 1~5w indicated the weight of rats on a normal/high-fat diet for 4 weeks, 5~9w indicated the weight of rats with different interventions for the following 4 weeks.

**Table 1.**
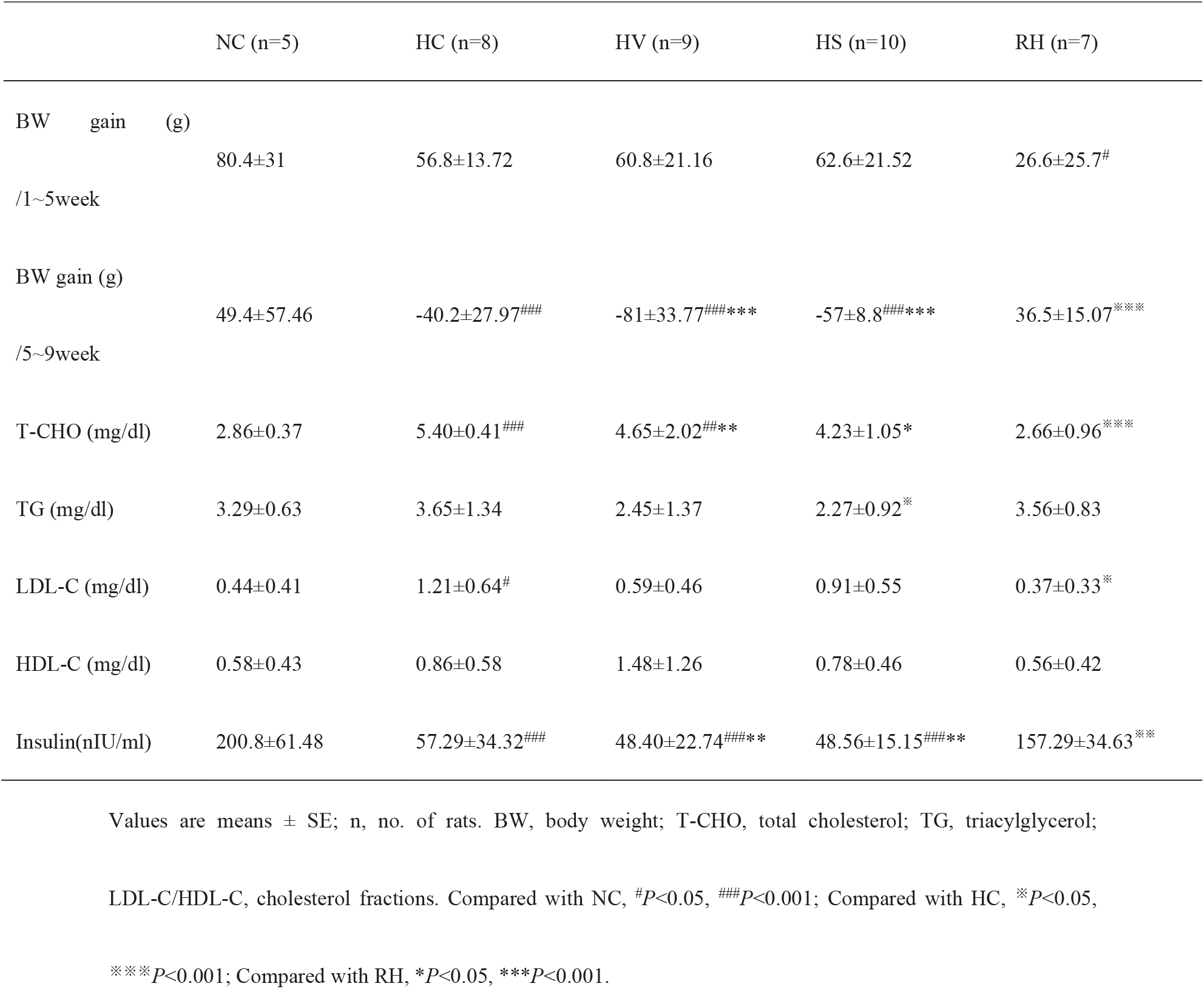
Changes in Body weight and Lipid metabolite concentrations

### 2 Morphological Changes of Adipose tissue on Oil Red O staining

The OD value of Oil Red O of different groups showed different histological changes on epididymis adipose (Table 2). Compared with NC, the OD value of HC and HV tended to be lower (*P*<0.05), but that in RH was higher (*P*<0.05). Compared with HC, higher OD value was shown in HS (*P*<0.05) and RH (*P*<0.01), but the OD value of RH was even higher than HS (*P*<0.05). According to Fig.3, normal lipid droplet in HC was not obvious among the high-fat diet rats, and most of them were partial flake with diffuse steatosis. HV group showed similar scattered lipid droplets as HC did. While it showed a lower degree of steatosis and some irregular red crystals could be ignored. Scattered lipid droplets were similar in HC and HV and the latter demonstrated a lower degree of steatosis. and some irregular red crystals could be ignored. There were also varies of steatosis in HS and RH, and the lipid droplets in these groups were more obvious than that of HC. The fat depot in RH was less punctate with little steatosis, while the amount of lipid droplets in the tissue was higher than that in HV and HS. Compared with other interventions, HS group showed similar normal lipid tissue in Oil Red O with NC, which means swimming was better to alleviate pathological adipose changes on high-fat diet rats.

**Fig.3.**
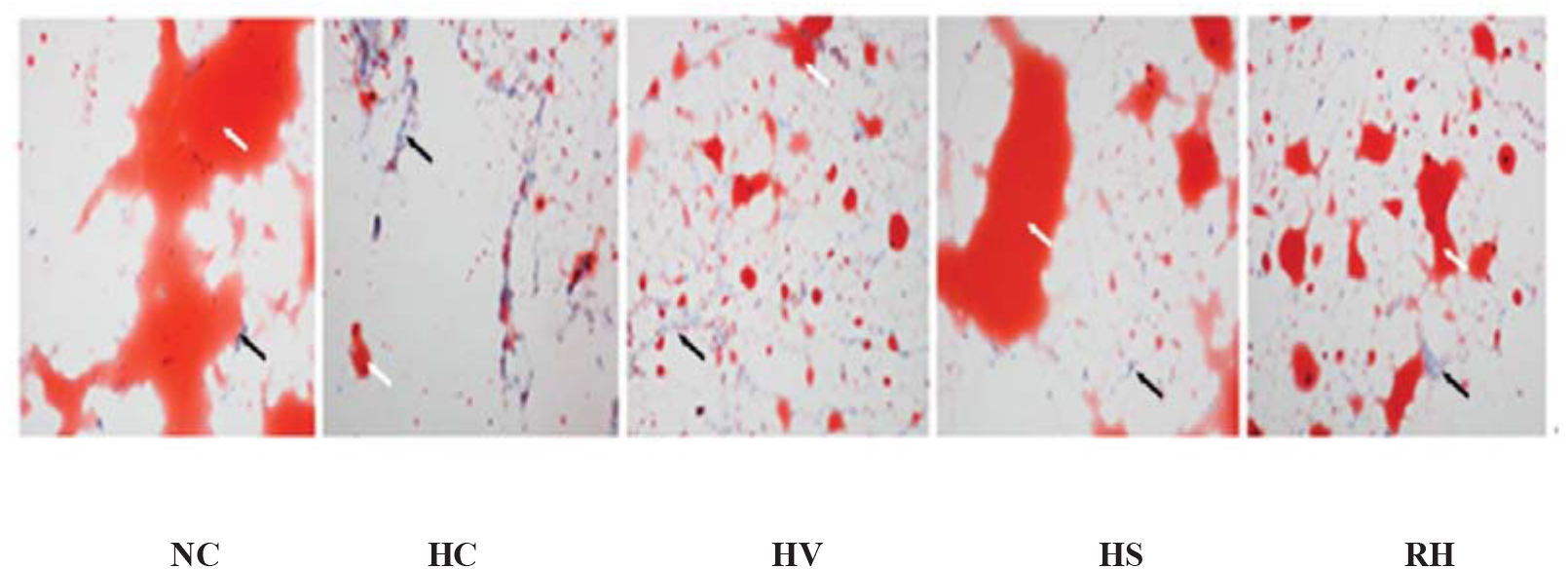
Oil Red O staining in Epididymis fat tissues of rats in each group (×200): note that, the white arrow showed normal lipid tissue which was stained with orange, and the black arrow showed steatosis undissolved in Oil Red O which nucleus was stained with blue.

**Table 2.**
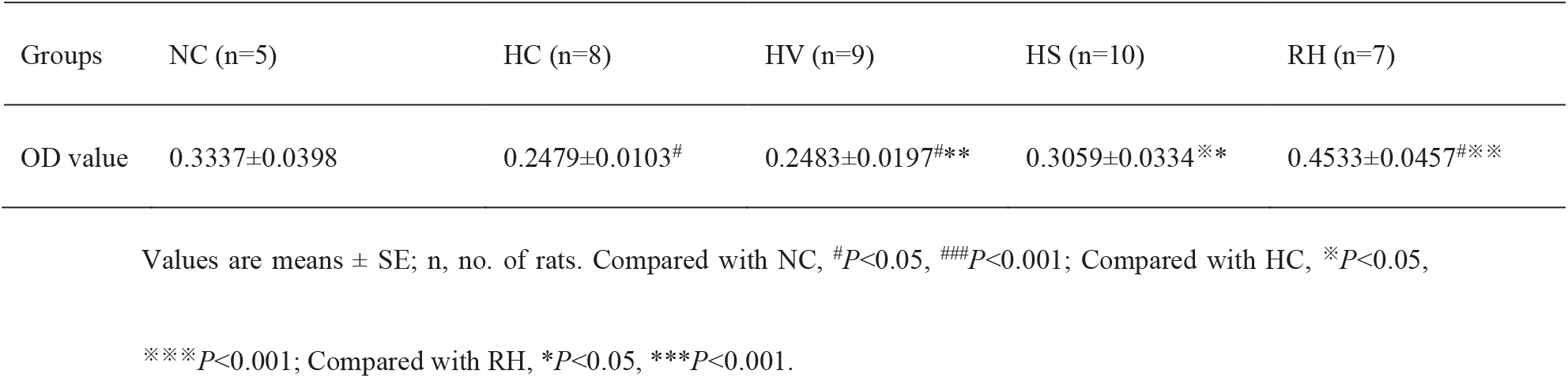
Changes in the OD value of Oil Red O on Epididymis adipose of rats

### 3 Serum Lipid and Insulin Levels from Blood

#### Serum lipid levels

As shown in Fig.2(C), the indexes of serum lipid (T-CHO, TG, LDL-C and HDL-C) in these groups were different, among which T-CHO was most obvious distinctly, TG and LDL-C were individual different, and HDL-C had no statistical difference. According to Tab.1, compared with NC, it was higher of T-CHO concentration in HC (*P*<0.001) and HV (*P*<0.01), and higher of LDL-C concentration in HC (*P*<0.05). Compared with HC, HS was significantly lower in TG (*P*<0.05), and RH was significantly lower in T-CHO (*P*<0.001) and LDL-C (*P*<0.05); Compared with RH, there was significant higher of T-CHO concentration in HV (*P*<0.01) and HS (*P*<0.05). Therefore, swimming could specifically reduce TG concentration. And refeeding diet could specifically reduce both T-CHO and LDL-C. While rice vinegar had no significant effect on improving blood lipid metabolism.

#### Serum insulin levels

It was shown significant difference in insulin levels among groups in Fig.2(D). Compared with NC, lower serum insulin level was observed significant in HC, HV and HS (*P*<0.001) in Tab.1, while there was no difference among these 3 groups. Specifically, the insulin levels in RH, which was higher than HC, HV and HS (*P*<0.01), did not show significant difference compared with NC. That is, the insulin level in high-fat diet rats was lower. It could be reversed by refeeding, rather than swimming or rice vinegar.

### 4 Expression of AMPKα in Pancreas, Liver and Cardiac tissues

As shown in Fig.4(A), AMPKα expression in pancreas of each group was significantly different. The expression in rice vinegar, swimming and refeeding groups was significantly higher than that of other groups (P<0.001). It was even higher in HS group than that in refeeding group (P<0.001). AMPKα expression of liver shown in Fig.4(B) indicated that there was no distinct difference among these groups. While different interventions did affect liver AMPKα which was higher in RH (*P*>0.05) and lower in HC (*P*>0.05). As shown in Fig.4(C), compared with NC group, cardiac AMPKα expression in each group increased after high-fat diet, and the expression was significantly higher in rice vinegar group (p<0.01) and swimming group (p<0.001). Therefore, different interventions had distinct effects on the expression of AMPKα in special tissues. Both swimming and rice vinegar had impacts on pancreas and cardiac tissues, while more obvious influence was shown on the swimming group. The refeeding group had a significant effect on the pancreas. The sign of its influence on liver tissue was spotted but no statistic difference was shown by far.

**Fig.4.**
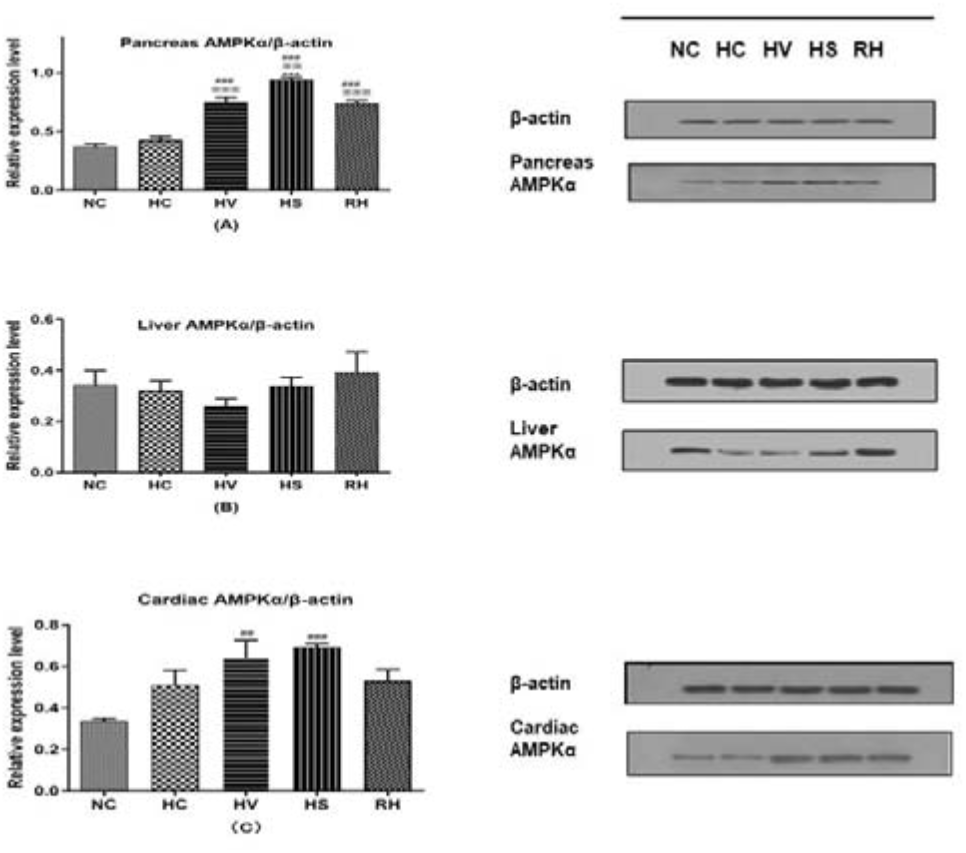
Relative expression of AMPKα in Pancreas, Liver and cardiac tissues. Values are means ± SE; n, no. of rats. Compared with NC, ^#^*P*<0.05, ^###^*P*<0.001; Compared with HC, ^※^*P*<0.05, ^※※※^*P*<0.001; Compared with RH, \**P*<0.05, \*\*\**P*<0.001.(A) shows the relative expression of AMPKα in pancreas, and (B) shows the relative expression of AMPKα in liver.

According to results above that AMPKα expression in pancreas was higher after different interventions, we analyzed the relationships between pancreas and lipometabolism furtherly. The correlation between pancreas AMPKα and blood lipid discussed subsequently by Pearson analysis in Tab.3. It turned out that, among all the lipid metabolite concentrations, only TG was found to be linked with pancreatic AMPKα (P<0.05). That is, swimming could specifically reduce the TG concentration from blood lipid and showed more effect on pancreas AMPKα than rice vinegar or refeeding did.

**Table 3.** Correlation coefficient between Blood lipid and Pancreas AMPKα in rats

| Pearson | Pancreas AMPKα expression | |
| --- | --- | --- |
|  | r | p |
| T-CHO | -0.101 | 0.691 |
| TG | -0.569 | 0.014* |
| LDL-C | -0.055 | 0.829 |
| HDL-C | 0.045 | 0.861 |

Values are means ± SE; \* $P$ <0.05.

### 5 Expression of HNF1α and CTRP6 in tissues

As shown in Fig.5(A) and Fig.5(B), there was no significant difference between NC and other 4 groups (*P*>0.05) on the expression of HNF1α in liver and CTRP6 in periepididymal adipose, nor the difference was found among HV, HS and RH (*P*>0.05). While various expression levels of HNF1α and CTRP6 on each group affected by different methods indicated distinct outcomes in Fig.4. Theoretically, effective interventions probably functioned with higher HNF1α level and lower CTRP6 compared with HC. But the results did not show obvious effects on high-fat diet rats with lower HNF1α expression by rice vinegar, swimming or refeeding. In addition, except for RH, CTRP6 expression did not turn to be lower in HV or HS. A further analysis showed there was neither significant relationship between liver HNF1α and AMPKα in different tissues nor correlation among blood lipid, pancreas AMPKα and CTRP6 protein in rats in Tab.4 and Tab.5.

**Fig.5.**
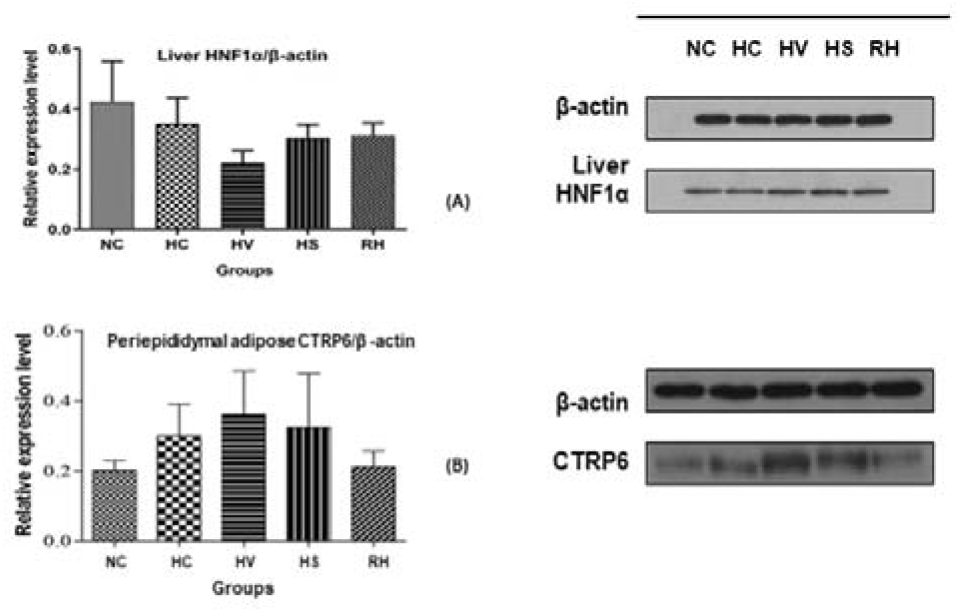
Relative expression of HNF1α in Liver and CTRP6 in Periepididymal adipose. Values are means ± SE; n, no. of rats. Compared with NC, ^#^*P*<0.05, ^###^*P*<0.001; Compared with HC, ^※^*P*<0.05, ^※※※^*P*<0.001; Compared with RH, \**P*<0.05, \*\*\**P*<0.001. (A) shows the relative expression of HNF1α in liver, and (B) shows the relative expression of CTRP6 in periepididymal adipose.

**Table 4.**
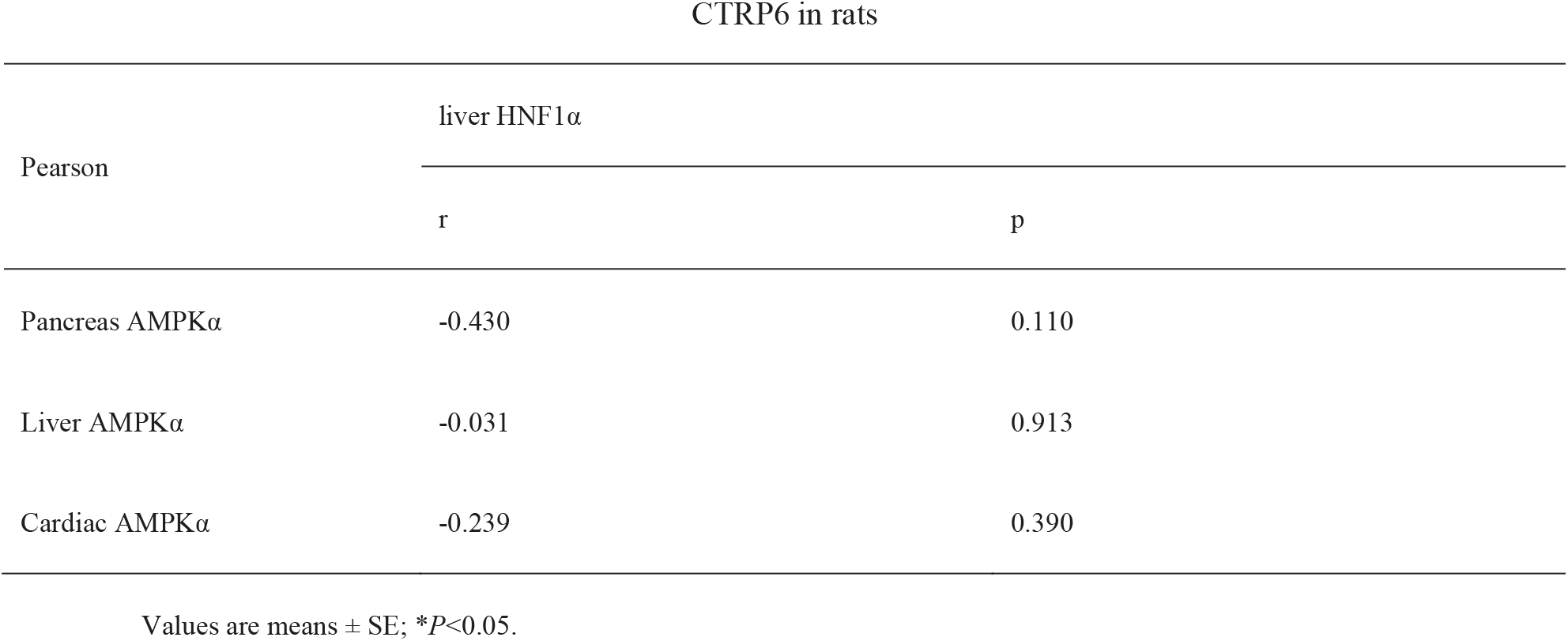
Correlation coefficient of blood lipid and Pancreas AMPKα with Periepididymal adipose CTRP6 in rats

**Table 5.**
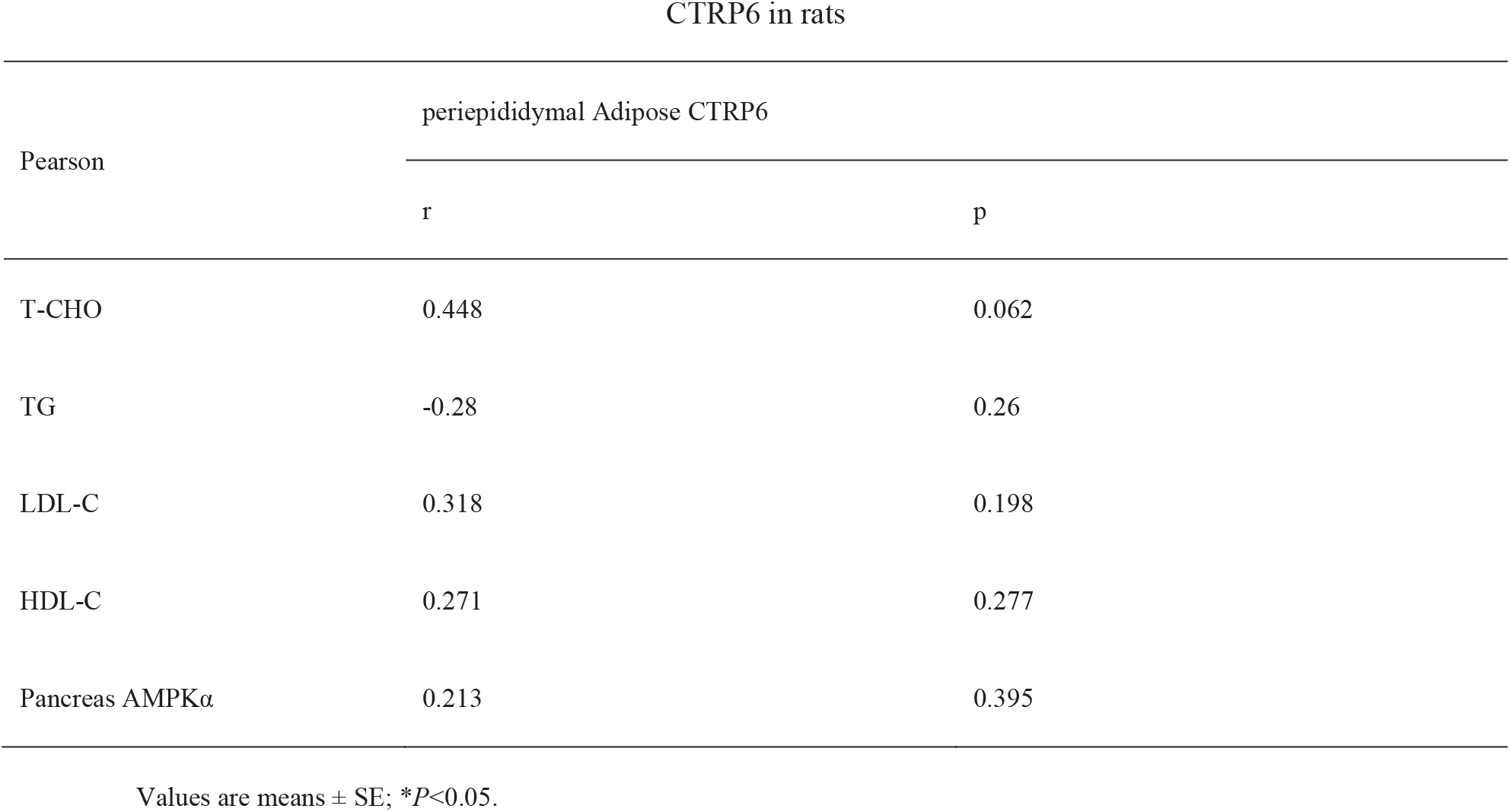
Correlation coefficient of blood lipid and Pancreas AMPKα with Periepididymal adipose CTRP6 in rats

## 6. Discussion

High-fat diet leads to abnormal lipometabolism and lipid homeostasis disorder, such as dyslipidaemia. Dyslipidaemia reflects the abnormal conditions of lipometabolism in vivo which mainly including T-CHO, TG and LDL-C. In this study we demonstrated different interventions by rice vinegar, swimming and refeeding on high-fat diet rats might lead to discrepant effects. Rice vinegar regulated the AMPKα expression of pancreas and cardiac tissues in high-fat diet rats, while it did not change obviously in blood lipid metabolism or steatosis. Swimming was a persuaded method to alleviate steatosis, along with losing weight, reducing TG concentration and insulin level. And its effect on regulating the indexes of serum lipid was related to the specific expression of AMPKα in pancreas. The results of refeeding showed, although there was no function of losing weight on refeeding rats, the content of T-CHO and LDL-C, insulin level and AMPKα expression in pancreas and liver tissues were regulated significantly. In general, in addition to rice vinegar group, the interventions of swimming and refeeding did play important role in regulating blood lipid metabolism. However, we also noticed that their effects on body weight and insulin level were completely opposite, and blood lipid composition was also different. The above conclusions provoke our further discussion: How do dyslipidemia, body weight and insulin level work on each other? And what is the possible pathway related to AMPKα and lipid metabolic regulation in various tissues?

Previous studies demonstrated that weight gain was a main manifestation on dyslipidaemia, especially on T-CHO, TG and lipoprotein in serum lipids (23). In our study, however, continuous high-fat diet had an effect of weight loss, which was even more obvious in rice vinegar and swimming groups. On the contrary, the rats of refeeding group gained weight obviously, which was closer to normal diet rats and had significant function on regulating serum lipid and insulin level. It gave us a second thought that body weight, though correlated to dyslipidaemia, might not be used as a direct indicator to reflect the level of lipometabolism. Compelling evidence supported the main effect by dyslipidemia was cardiovascular and fatty liver diseases particularly on the increase of T-CHO and LDL-C (8, 24). Refeeding diet, superior to swimming or rice vinegar, significantly helped to regulate dyslipidemia specifically by lowering T-CHO and LDL-C in our experiment. They were consistent with the previous conclusion that the effect of diet was mainly lowering TG and LDL-C, and exercise was regulating predominantly by lowering fasting TG (25). Dyslipidemia along with higher storage of TG by dietary intake probably caused abnormal insulin level which in turn impacted exogenous lipid by either inhibiting synthesis or promoting catabolism of TG (26, 27). We also concluded that rats with high-fat diet showed significant decrease of insulin level in serum which increased more obviously in refeeding rats than that of other interventions. Thus, it was suggested that insulin level probably regulated dyslipidaemia with balanced TG level, but whether it could restrict the content of TG continuously needs further study.

Studies on dietary restrictions have shown that altered meal pattern could be used to prevent certain metabolic diseases(28). Altered dietary may stimulate the mechanism against weight gain, fat accumulation, inflammation, glucose intolerance, insulin resistance, and loss of endurance and motor coordination(29). Our study reveals that refeeding from high-fat diet improves lipid homeostasis, which is seldom discussed by other researches. Specifically, the regained body weight and lower T-CHO and LDL-C by refeeding from high-fat diet was followed by significantly higher of insulin levels, which is an important marker of regulating lipometabolism globally. In addition, the fat depot in RH with less punctate steatosis means histological changes of lipid droplets was recovered partly by refeeding from high-fat diet. AMPKα expression of RH was significantly higher in pancreas and liver than high-fat diet rats. HNF1α is a typical protein of liver. The expression of it in RH was higher than that in HV or HS but still lower than HC. On the contrary, CTRP6 of periepididymal adipose in RH was closer to that in normal diet group. Above results highlight therapeutic potentials of refeeding diet, independent of alterations in weight loss or other consumptions, as a persuaded advice to ameliorate dyslipidaemia changes in lipometabolic-associated pathology.

AMPKα is regarded as a target of low energy regulation and unstable insulin level affected by abnormal lipometabolism, such as synthesis of fatty acids and cholesterol (30, 31). In order to regulate lower expression of AMPKα, rice vinegar, swimming and refeeding were intervened to these high-fat diet rats. Our study showed significantly higher AMPKα expression particularly in pancreas among different tissues after intervened high-fat diet rats. And it showed a significant negative correlation between AMPKα in pancreatic and TG. As it was discussed in our study that rats intervened only by swimming showed lower TG and higher expression of pancreas AMPKα. We got the hypothesis that swimming, rather than refeeding, could affect dyslipidaemia by specifically moderating pancreas AMPKα.

The intervention of refeeding showed a superior effect on liver AMPKα and insulin level than others. The probably mechanism of it on insulin level and lipometabolic disorder might be related to targeted tissues of insulin, i.e., liver and surrounding adipose. HNF1α and CTRP6 are supposed to be the substance metabolic indicators of these two targets during lowering lipid (32, 33). Deficiency of HNF1α in liver lipometabolism may lead to diabetes and hypercholesterolemia by increasing bile acid and cholesterol synthesis (34, 35). The cross-talking of insulin level and AMPKα/HNF1α supports AMPK-mTOR-HNF1α pathway for the treatment of lipometabolism and insulin dysfunction (36). However, HNF1α expression in RH did not show lower than that in HC, which means the effect on dyslipidaemia by refeeding might not work through AMPK-mTOR-HNF1α signal pathway. CTRP6, the main protein of periepididymal adipose, contains a globular domain which inhibits to mediate the phosphorylation and activation of muscular AMPK. Researches have discussed its function of participating in fatty acid metabolism and proliferation of ectopic adipocytes through AMPK/ACC pathway (32, 37) (38). Our study showed the expression of CTRP6 in adipose tissue was only decreased in refeeding rats, and its effect on lowering lipid may relate to liver AMPKα and periepididymal adipose CTRP6. Combined with significant changes of T-CHO in serum lipid and insulin level, the peripheral regulation on lipometabolism by refeeding probably mediates targets from liver to periepididymal. We will focus on AMPK-ACC-CTRP6 signal pathway for further research on this regulation in the subsequent experiments.

This study demonstrates a preferred method of ameliorating dyslipidemia by refeeding which is better than swimming or rice vinegar on high-fat diet rats. There is a noteworthy peripheral regulation on lipometabolism by refeeding normal diet. We supposed it is mainly triggered by targeted tissues from liver to periepididymal, and then resulted in the significant changes of serum lipid T-CHO and insulin level. While in the process of this experiment, there are some defects as following: Analysis of food intake should be added in the experiment, or the rats should be given a fixed diet intake to be compared among different groups; The decline in lowering lipid was accompanied by peripherally regulating AMPKα, CTRP6, and insulin levels. Future works should be strived to understand roles of these indexes in some pathways, such as AMPK-ACC-CTRP6; It is limited study for the lipid lowering mechanism of rice vinegar and swimming, which might function on specific proteins, such as pancreas AMPKα and related pathway.

In summary, refeeding and swimming significantly regulate lipometabolic related indicators after high-fat diet through different target organs, such as AMPKα in pancreas, liver and cardiac tissues. Refeeding diet, compared with other interventions, functioned better in regulating the lipometabolic level after high-fat diet. Whatever approach mentioned above we adopted to intervene, the best policy to keep the balance of lipid homeostasis is to maintain a healthy diet.

## Abbreviations

AMPK: AMP-activated protein kinase
CTRP: C1q/TNF-associated protein
HNF1α: hepatocyte nuclear factor-1 α
HDL: high density liptein
LDL: low density liptein
TG: triacylglycerol
T-CHO: total cholesterol
ACC: acetyl CoA carboxylase
FA: fatty acids
Glu: glucide
NC: normal diet group
HC: high-fat control group
HV: high-fat diet with rice vinegar group
HS: high-fat diet with swimming group
RH: refeeding normal diet group
BW: body weight

## 7. Acknowledgments/grant support

## Author Contributions

Author contributions: YY conception and design of research; ZF performed experiments; YY, ZF analyzed data; YY, ZF interpreted results of experiments; YY prepared figures; YY, ZF drafted manuscript; LH, MC, XX edited and revised manuscript; YY, LH approved final version of manuscript.

